# Human caspase-4 and caspase-5 assemble into non-canonical inflammasomes in response to overlapping and distinct LPS structures

**DOI:** 10.64898/2026.10.06.756901

**Authors:** Ling Wu, Kimberly A. Wodzanowski, Stephanie M. Schreiner, Klarissa I. Diaz, Katharine B. Fox, Justin Su, Michael Kegel, Srikar Yelamarthy, Jenna Muscat-Rivera, Daniel Grubaugh, Madhura Kulkarni, Mallory A. Evans, Ricardo O. Carale, Matthew E. Sherman, Cornelius Y. Taabazuing, Robert K. Ernst, Jilian Melamed, Edward F. Kreider, Igor E. Brodsky, Sunny Shin

## Abstract

Innate immune sensing of bacterial lipopolysaccharide (LPS) is critical to host defense, but a dysregulated response can lead to sepsis, a life-threatening condition associated with multiple organ failure and shock. LPS is sensed at the cell surface by Toll-like receptor 4 (TLR4) and in the cytosol by mouse caspase-11 (Casp11) or its human orthologs, caspase-4 and 5 (CASP4/5). In response to cytosolic LPS, Casp11 oligomerizes in a catalytically dependent manner into supramolecular organizing centers (SMOCs). We demonstrate that CASP4 and CASP5 also form catalytically dependent SMOCs in response to bacterial LPS in macrophages. Furthermore, we find that CASP4 and CASP5 differentially respond to distinct LPS structural variants that fail to activate CASP11 and that only activating LPS structures induce SMOC formation by the corresponding caspase. Notably, transgenic mice expressing CASP4 in lieu of Casp11 exert more effective control of *Francisella novicida*, a rodent pathogen whose LPS evades TLR4 and Casp11 detection. Conversely, CASP4-expressing mice are more susceptible to sepsis when exposed to *E. coli* expressing an LPS variant that fails to activate TLR4 and Casp11. Together, these findings reveal human CASP4 and CASP5 have evolved to assemble into SMOCs in response to a broader range of LPS structures than mouse Casp11 and highlight the potential tradeoff between systemic inflammation and control of bacterial infection that may underline these differences. Our findings may inform the development of targeted therapeutic strategies for gram-negative bacterial infection and sepsis that are effective in humans.

## INTRODUCTION

The mammalian immune system relies on a highly conserved set of pattern recognition receptors (PRRs) to detect pathogen-associated molecular patterns (PAMPs) present on gram-negative bacteria, mount appropriate immune responses, and clear invading pathogens. Specifically, the myeloid differentiation 2/Toll-like receptor 4 (MD-2/TLR4) complex detects lipopolysaccharide (LPS), a key defining characteristic of gram-negative bacterial cell walls, at the cell surface, while the murine cysteine protease caspase-11 (Casp11) senses LPS within the cytosol of host cells^1–3^. Casp11 subsequently becomes active and oligomerizes into the noncanonical inflammasome. Active Casp11 cleaves and activates the host protein gasdermin-D (GSDMD), which forms plasma membrane pores, allowing for release of IL-1 family cytokines and pyroptotic cell death. In murine models, Casp11 is a key driver of both host defense and septic shock in addition to TLR4^1,2^.

Humans possess two Casp11 orthologs, caspase-4 (CASP4) and caspase-5 (CASP5), that also sense cytosolic LPS. Like Casp11, CASP4/5 activation leads to downstream GSDMD cleavage and the release of IL-1 cytokines^4–6^. CASP4/5/11 are composed of the N-terminal caspase activation and recruitment domain (CARD) and the catalytic domain, which is comprised of the p20 and p10 subunits connected by the interdomain linker (IDL)^7^. LPS binding directly to the CARD domain of caspase 4/5/11 initiates caspase autoprocessing and downstream non-canonical inflammasome activation. However, the molecular mechanisms by which LPS promote caspase-4/5/11 inflammasome formation remain poorly understood. We have shown previously that in response to cytosolic LPS, Casp11 assembles into oligomeric structures within the cell that rely on Casp11 enzymatic activity^8^. Whether and how CASP4/5 also form oligomeric structures in response to cytosolic LPS has not been investigated.

We and others have previously shown that CASP4/5 mediate inflammasome responses to gram-negative bacteria and LPS in human cells^9–12^, but it is unclear whether CASP4 and CASP5 have distinct or redundant roles. Several bacterial pathogens evade innate immune sensing by modulating their LPS structure. *Yersinia pestis* specifically modifies its LPS acylation state from hexa-acylated LPS at 26°C that robustly activates TLR4 and CASP11 to its non-activating tetra-acylated form at 37°C^1,13,14^. Intriguingly, studies from our labs and others indicate that human CASP4/5 respond to a broader range of LPS structures than Casp11 or TLR4, but whether CASP4 and CASP5 respond to overlapping or distinct LPS structures is unclear^10,11,15^. Furthermore, both activating and non-activating LPS structures are reported to bind Casp11, CASP4, and CASP5^4^, raising questions about whether specificity for distinct LPS structures is determined by downstream steps such as LPS-induced oligomerization.

Here, we report that like Casp11, CASP4 and CASP5 form SMOCs in response to intracellular LPS. As expected, CASP4 and CASP5 both activate the inflammasome response against hexa-acylated LPS. Interestingly, we found that CASP4 and CASP5 exhibited specificity for distinct LPS structural variants that do not activate Casp11. CASP4 underwent oligomerization and activation in response to a tetra-acylated LPS variant, but not to a penta-acylated LPS variant. In contrast, CASP5 underwent oligomerization and activation in response to a penta-acylated LPS variant, but not to the tetra-acylated LPS variant. Lastly, using *Casp11*^-/-^ transgenic mice expressing *CASP4*, we observe that CASP4 provided protection against tetra-acylated LPS-expressing *Francisella novicida,* but also induced septic shock in response to *E. coli* expressing tetra-acylated LPS. Together, these data suggest that CASP4/5 play both complementary and distinct roles and have evolved to detect a variety of LPS structures to maximize pathogen detection.

## RESULTS

### Spontaneous CASP4 and CASP5 speck formation in HEK293T cells is dependent on their catalytic activity

Canonical inflammasomes typically consist of a sensor protein belonging to the NLR family, the adaptor protein ASC, and caspase-1. Upon activation, the canonical inflammasome forms SMOCs that can be visualized under the microscope as a large speck^16–18^. Recently, we demonstrated that mouse Casp11 also forms SMOCs during inflammasome activation despite lacking an upstream sensor or adaptor^8^. To date, whether human CASP4 and CASP5 form SMOCs upon LPS stimulation has remained elusive. Assessing endogenous CASP4 and CASP5 speck formation via microscopy have been limited by the lack of endogenous CASP4- and CASP5-specific antibodies suitable for immunofluorescence. We therefore initially employed the same approach we previously used to study Casp11 oligomerization by fusing the mCherry tag to the C-terminus of CASP4 and 5 (**Figure S1A**). In HEK293T cells, which normally do not express inflammasome components, ectopic expression of plasmids encoding Casp11 results in spontaneous speck formation and caspase activity^8,19–22^. Surprisingly, we found that C-terminally-tagged CASP4 and CASP5 had poor speck formation, in contrast to Casp11, and induced minimal cytotoxicity when co-expressed with GSDMD (**Figure S1B**). Moreover, C-terminally-tagged CASP4 and CASP5 showed limited cleavage of IL-18 and minimal release of cleaved IL-18 into the supernatant (**Figure S1C**), suggesting that unlike with Casp11, appending a C-terminal tag to CASP4 or CASP5 interfered with their activity. We therefore moved the mCherry tag to the N-terminus of CASP4 (mCh-CASP4) or CASP5 (mCh-CASP5) (**Figure 1A**), which resulted in robust GSDMD and IL-18 processing and cell death (**Figure 1B, E-F, S1C-D**), suggesting that N-terminally-tagged CASP4 and CASP5 remain viable enzymes. Furthermore, ectopic expression of mCh-CASP4 or mCh-CASP5 in HEK293T cells led to spontaneous speck formation (**Figure 1C-D**) that was not observed with C-terminally tagged CASP4 or CASP5 (**Figure S1B**). Moreover, this spontaneous speck formation was abrogated by treatment with the pan-caspase inhibitor zVAD, (**Figure 1C-D**), consistent with our prior studies demonstrating that spontaneous formation of Casp11 specks under these conditions requires its catalytic activity^8^. Consistently, mutation of the catalytic cysteine to an alanine in CASP4 (C258A) and CASP5 (C315A) abrogated both their ability to cleave substrates GSDMD and IL-18, and to form spontaneous specks in HEK293T cells (**Figure 1E-H**). These findings demonstrate that, like Casp11, the catalytic activity of CASP4 and CASP5 is required for their oligomerization and speck formation.

**Figure 1:**
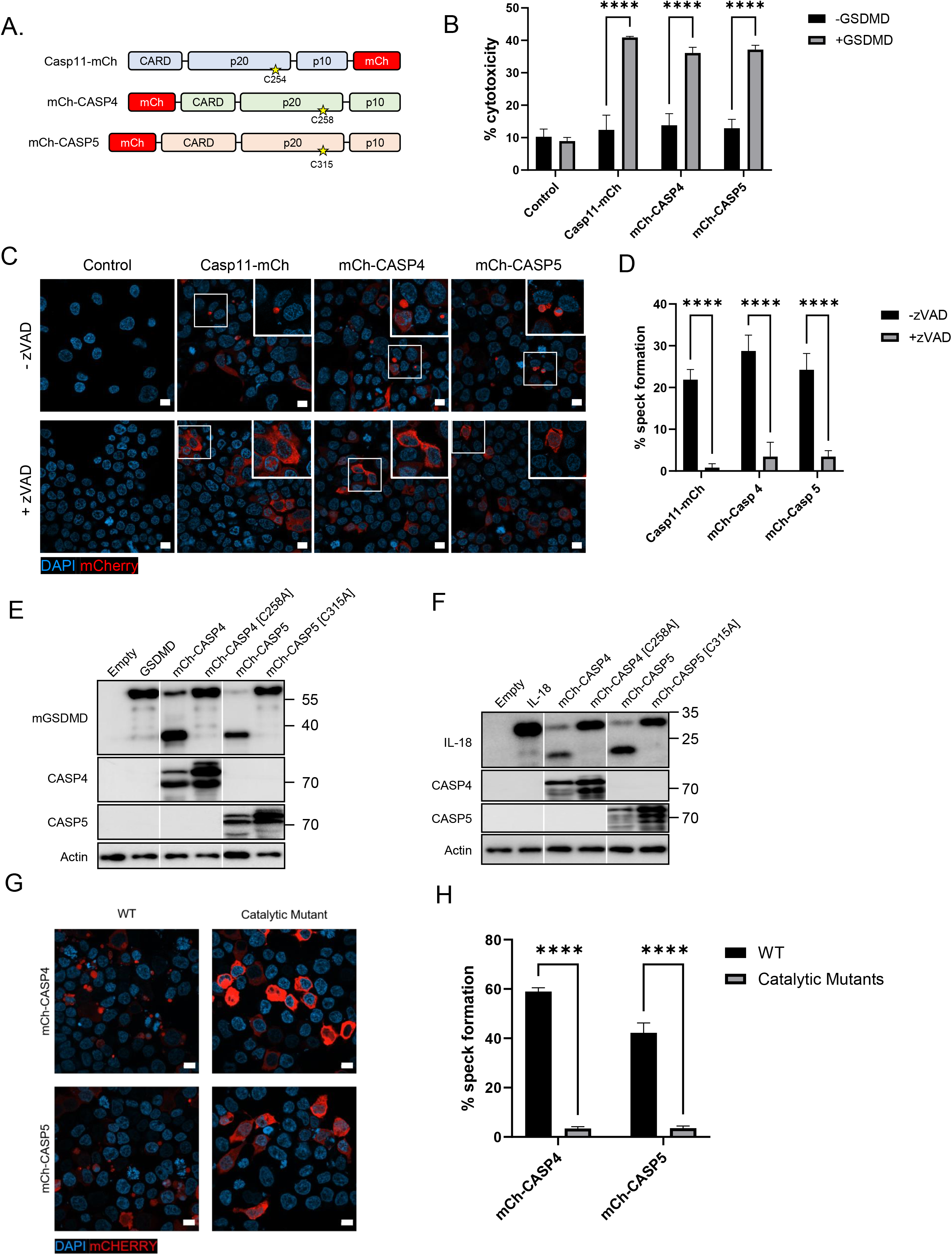
Human CASP4 and CASP5 form spontaneous and catalytically dependent functional specks in HEK293T cells. (A) Schematic representation of mCherry (mCh)-tagged Casp11, CASP4, and CASP5. The catalytic cysteine residues are indicated by the yellow star for each caspase. (B) HEK293T cells were transfected with plasmids encoding the indicated mCh-tagged caspase and GSDMD. Cytotoxicity was measured based on lactate dehydrogenase (LDH) release at 24 hours transfection. Shown are pooled data of 3 independent biological replicates. (C) HEK293T cells transfected with indicated mCh-tagged constructs with or without Z-VAD (200 µM) for 24 hours and visualized by fluorescence microscopy. Nuclei were stained with DAPI. Scale bar = 10µm. (D) Speck formation in (C) was quantified as the percentage of mCh-CASP4- or mCh-CASP5-expressing cells containing at least one speck. Pooled data of 3 biological replicates with 150-200 cells analyzed per condition ****p<0.0001, Two-way ANOVA with Sidak’s multiple-comparison test. (E-F) HEK293T cells transfected with mCh-tagged caspase, GSDMD, and IL-18 (125ng each plasmid). Immunoblot analysis at 24 hours post-transfection for CASP4, CASP5, GSDMD, and IL-18 as indicated. β-actin was used as a loading control. Representative of at least 3 independent biological replicates. (G) Fluorescence microscopy of HEK293T cells transfected with indicated mCh-tagged constructs (200 µM) for 24 hours. Nuclei were stained with DAPI. Scale bar = 10µm. (H) Speck formation in (G) was quantified as the percentage of mCh-CASP4- or mCh-CASP5- or catalytic mutant-expressing cells containing at least one speck. Pooled data of 3 biological replicates with 150-200 cells analyzed per condition. ****p<0.0001, Two-way ANOVA with Sidak’s multiple-comparison test.

### Cytosolic LPS induces CASP4 and CASP5 speck formation in macrophages

To develop a more physiological model where we could test whether LPS can induce CASP4 or CASP5 speck formation in macrophages, we generated HOXB8-conditionally immortalized *Casp11^-/-^* bone marrow progenitors^23^. This cell background eliminates endogenous Casp11 activity and permits the stable expression of mCh-CASP4 or mCh-CASP5 following lentiviral transduction and selection. Following differentiation into macrophages, these immortalized bone marrow-derived macrophages (iBMDMs) exhibited robust expression of mCh-CASP4 or mCh-CASP5 (**Figure 2A**). Furthermore, the mCh-CASP4- or mCh-CASP5-expressing macrophages underwent cell death in response to LPS transfection (**Figure 2B**). Critically, we observed that both mCh-CASP4 and mCh-CASP5 formed LPS-inducible specks at six hours post-transfection with *E. coli* LPS (**Figure 2C-D**). Moreover, infection with the intracellular gram-negative bacterium *Legionella pneumophila* likewise induced mCh-CASP4 or mCh-CASP5 specks (**Figure 2E-F**). These data indicate that CASP4 and CASP5 undergo oligomerization in response to both cytosolic LPS and gram-negative bacterial infection.

**Figure 2.**
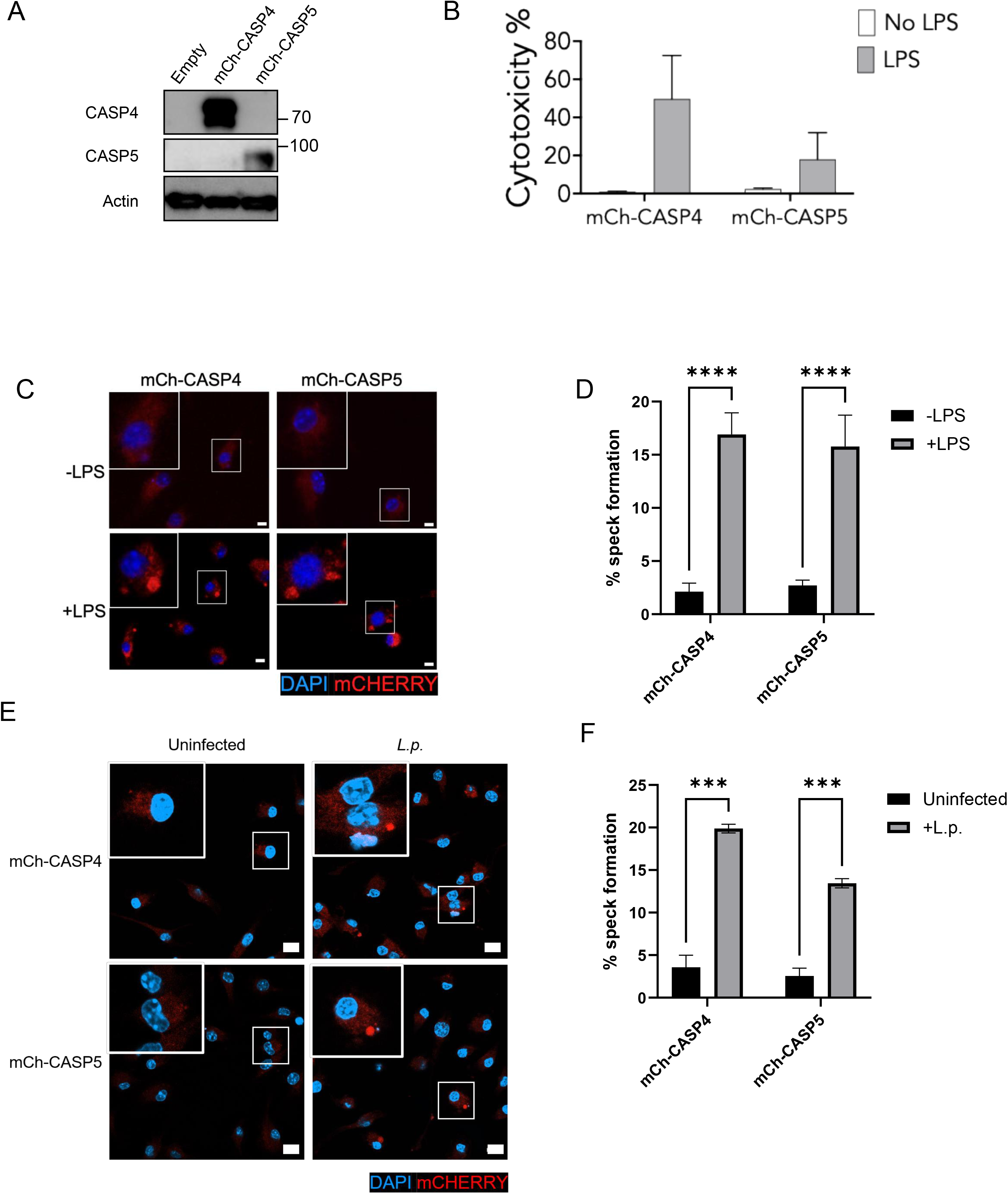
CASP4 and CASP5 assemble into oligomeric specks in response to cytosolic LPS in macrophages. (A) *Casp11*^-/-^ immortalized BMDMs (iBMDMs) were transduced with lentivirus encoding mCherry-CASP4 (mCh-CASP4) or mCherry-CASP5 (mCh-CASP5) and assessed via immunoblot analysis to confirm caspase expression. (B) mCh-CASP4- or mCh-CASP5-expressing iBMDMs were transfected with *E. coli* LPS and assessed for cytotoxicity at 16 hours following LPS stimulation. Results show pooled data from two independent experiments. (C) Cells were transfected with *E. coli* LPS and analyzed by fluorescence microscopy 6 hours following transfection. Nuclei were stained with DAPI. Scale bar = 10µm. (D) Speck formation in (C) was quantified as the percentage of mCh-CASP4- or mCh-CASP5-expressing cells containing at least one speck. Pooled data of 3 biological replicates with 150-200 cells analyzed per condition. ***p<0.001, Two-way ANOVA with Sidak’s multiple-comparison test. (E) mCh-CASP4- or mCh-CASP5-expressing iBMDMs were left uninfected or infected with *L. pneumophila* (MOI=10) for 6 hours followed by fluorescence microscopy analysis for speck formation (F). Nuclei were stained with DAPI. Pooled data of 2 biological replicates with 150-200 cells analyzed per condition ****p<0.0001, Two-way ANOVA with Sidak’s multiple-comparison test.

### Casp11, CASP4, and CASP5 respond to overlapping and distinct LPS structures

Many gram-negative bacterial pathogens modify their LPS structures to evade immune detection^24^. One of the best-characterized LPS modifications is a change in lipid A acylation state^14,25^. Prototypical hexa-acylated lipid A (**Figure 3A**) robustly activates TLR4 and both mouse and human non-canonical inflammasomes, whereas under-acylated lipid A structures, such as penta- and tetra-acylated lipid A (**Figure 3A**), elicit more variable immune responses^10,13,15^. We and others have found that while hexa-acylated lipid A robustly activates both human CASP4 and murine Casp11, the tetra-acylated lipid A structures found in immunoevasive pathogens like *Yersinia pestis* or *Francisella novicida* robustly activate CASP4 but fail to activate mouse and human TLR4 or Casp11^10,11,15^. We set out to recapitulate these findings by ectopically expressing Casp11 or CASP4 in murine BMDMs. As expected, bone marrow-derived macrophages (BMDMs) from WT mice responded robustly to transfected hexa-acylated LPS but not tetra-acylated LPS, as shown by robust GSDMD cleavage and high cytotoxicity (**Figure 3B-C**). BMDMs from *Casp11^-/-^* mice reconstituted with *Casp11* or *CASP4* exhibited GSDMD cleavage in response to hexa-acylated LPS transfection, whereas transfection of tetra-acylated LPS induced GSDMD cleavage and cytotoxicity only in cells reconstituted with *CASP4* but not *Casp11* (**Figure 3B-C**). These responses relied on Casp11 or CASP4, as *Casp11^-/-^* BMDMs transduced with empty vector failed to undergo GSDMD and cell death in response to LPS transfection (**Figure 3B-C**).

**Figure 3.**
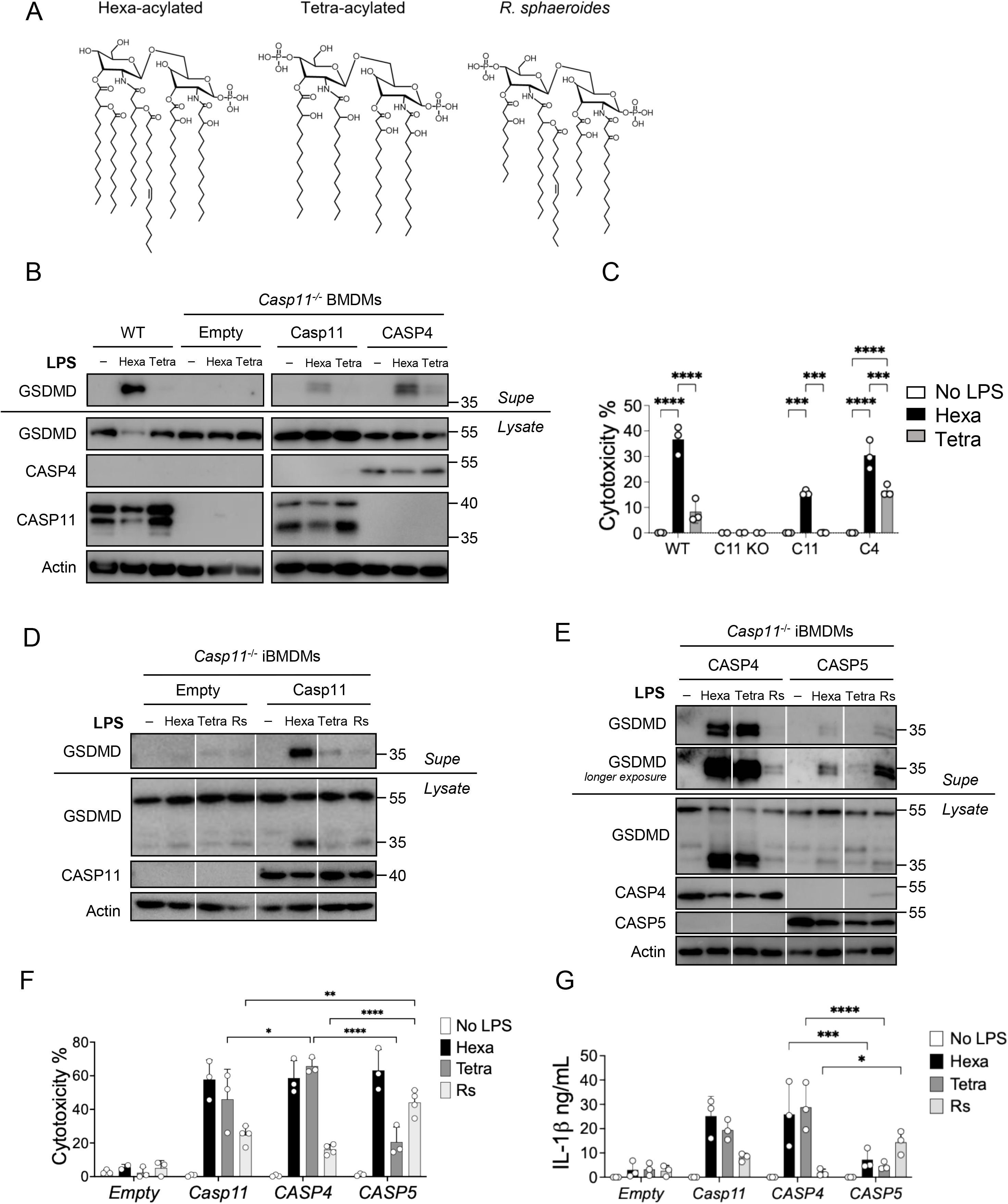
Casp11, CASP4, and CASP5 exhibit differential responses to distinct LPS structures. (A) Schematic diagram of LPS structures. Hexa-acylated (Hexa) and tetra-acylated (Tetra) LPS structures were isolated from *Yersinia pestis.* Penta-acylated structure was isolated from *Rhodobacter sphaeroides* (Rs). (B-C) Primary WT or *Casp11^-/-^*BMDMs transduced with lentivirus encoding *empty*, *Casp11*, or *CASP4* were primed with Pam3CSK4 (400ng/mL) for 4 hours, followed by LPS (2µg/mL) transfection. 16 hours after transfection, lysates and TCA-precipitated supernatants were immunoblotted for the indicated proteins (B). Supernatant were assessed for cytotoxicity by LDH release assay (C). (D-E) *Casp11^-/-^*iBMDMs were reconstituted with 1µg/E6 cells of empty or LNPs encoding *Casp11*, *CASP4*, or *CASP5* for 48hrs. They were then primed with Pam3CSK4 (400ng/mL) for 4 hours, followed by intracellular stimulation with indicated LPS structures (2µg/mL) for 16hrs, and analyzed for the indicated proteins in the cell lysate or TCA precipitated supernatant by immunoblot, (D) cytotoxicity by LDH release assay, and (E) released IL-1*_β_* levels by ELISA. Results are representative of three independent experiments (B-C), or show pooled data from at least three independent experiments, with each dot representing the average of biological replicates from each experiment (D-G). n.s=not significant, *p≤0.05, **p≤0.01, ***p≤ 0.001, ****p≤ 0.0001 by two-way ANOVA.

Whether human CASP4 and CASP5 mount distinct or overlapping responses to different LPS structural variants is unclear. Because human macrophages express differing levels of CASP4 and CASP5, it has been challenging to define the ability of each caspase to sense distinct LPS structural variants. To address this problem, we applied the nucleoside-modified mRNA-lipid nanoparticle (mRNA-LNP) platform, which enables efficient and titratable ectopic protein expression in various cell types^26,27^.We generated mRNA-LNPs encoding *Casp11*, *CASP4*, and *CASP5* to permit functional dissection of each caspase’s role in responding to different LPS variants. Importantly, *Casp11^-/-^* macrophages individually expressing Casp11, CASP4, or CASP5 all cleaved mouse GSDMD in response to transfected *E. coli* hexa-acylated LPS (**Figure S2A**) and exhibited robust cytotoxicity and IL-1β release (**Figure S2B-C**). These findings indicate that the mRNA-LNP platform enables tunable caspase expression in macrophages in a manner that maintains their functional activity in response to cytosolic LPS.

We next ectopically expressed Casp11, CASP4, and CASP5 in murine macrophages in order to directly compare individual responses of these caspases to three distinct LPS structures. We employed: 1) hexa-acylated LPS, which activates CASP4 and Casp11, and is presumed to activate CASP5, 2) tetra-acylated LPS, which activates CASP4 but not Casp11, with unknown effects on CASP5, and 3) *Rhodobacter sphaeroides* LPS (Rs-LPS), a well-described penta-acylated LPS antagonist that does not activate Casp11 or CASP4^4^, with unknown effects on CASP5. As expected, cytosolic hexa-acylated LPS induced GSDMD cleavage in *Casp11^-/-^* iBMDMs reconstituted with Casp11, CASP4, or CASP5 (**Figure 3D-E**). Tetra-acylated LPS, on the other hand, led to robust GSDMD cleavage in cells reconstituted with CASP4, but not those reconstituted with Casp11, in agreement with our and others’ previous data that human and mouse non-canonical inflammasomes differentially respond to tetra-acylated LPS^10,15^. Interestingly, unlike cells expressing CASP4, cells reconstituted with CASP5 did not undergo GSDMD cleavage in response to tetra-acylated LPS, suggesting that the human-specific response to tetra-acylated LPS is mediated primarily through CASP4. Consistent with the literature that Rs-LPS is a non-activating LPS, neither mouse Casp11-nor human CASP4-reconstituted cells exhibited GSDMD cleavage when stimulated with cytosolic Rs-LPS. Surprisingly, iBMDMs reconstituted with CASP5 exhibited GSDMD cleavage upon stimulation with cytosolic Rs-LPS. Notably, the presence of cleaved GSDMD in response to the different LPS variants correlated with cytotoxicity and IL-1β release (**Figure 3F-G**). Altogether, these data indicate that Casp11, CASP4, and CASP5 generate inflammasome responses to both unique and shared LPS structures.

### CASP4- and CASP5-specific responses to LPS variants are recapitulated in human macrophages and are dose-dependent

We next asked whether the differential responses of CASP4 and CASP5 to distinct LPS variants could be recapitulated in human macrophages. We reconstituted *CASP4^-/-^CASP5^-/-^*double-knockout (DKO) THP-1 macrophages with *CASP4-* or *CASP5*-encoding mRNA-LNPs followed by cytosolic LPS stimulation. Indeed, THP-1 cells reconstituted with CASP4 responded to hexa- and tetra-acylated LPS but not Rs-LPS, while CASP5-reconstituted THP-1 cells responded to hexa-acylated and Rs-LPS but not tetra-acylated LPS, as measured by cell death and IL-1β release (**Figure 4A-B**). To test whether increased expression levels of CASP4 or CASP5 would enable cells to respond to noncognate LPS variants, we reconstituted THP1 cells with increasing concentrations of *CASP4* or *CASP5* LNPs (**Figure 4C-F**) and stimulated the reconstituted cells with each of the three distinct LPS structures. THP-1 cells treated with increasing doses of *CASP4* LNPs displayed a corresponding dose-dependent increase in cell death in response to hexa- and tetra-acylated LPS, but not Rs-LPS (**Figure 4E**). In contrast, THP-1 cells treated with increasing doses of *CASP5* LNPs showed a dose-dependent increase in cell death in response to both hexa-acylated and Rs-LPS, but not tetra-acylated LPS (**Figure 4F**). Collectively, these data indicate that CASP4 and CASP5 respond to both overlapping and distinct LPS structures and reveal a unique ability of CASP5 to respond to an LPS variant not previously described to activate any other known mammalian LPS sensor.

**Figure 4.**
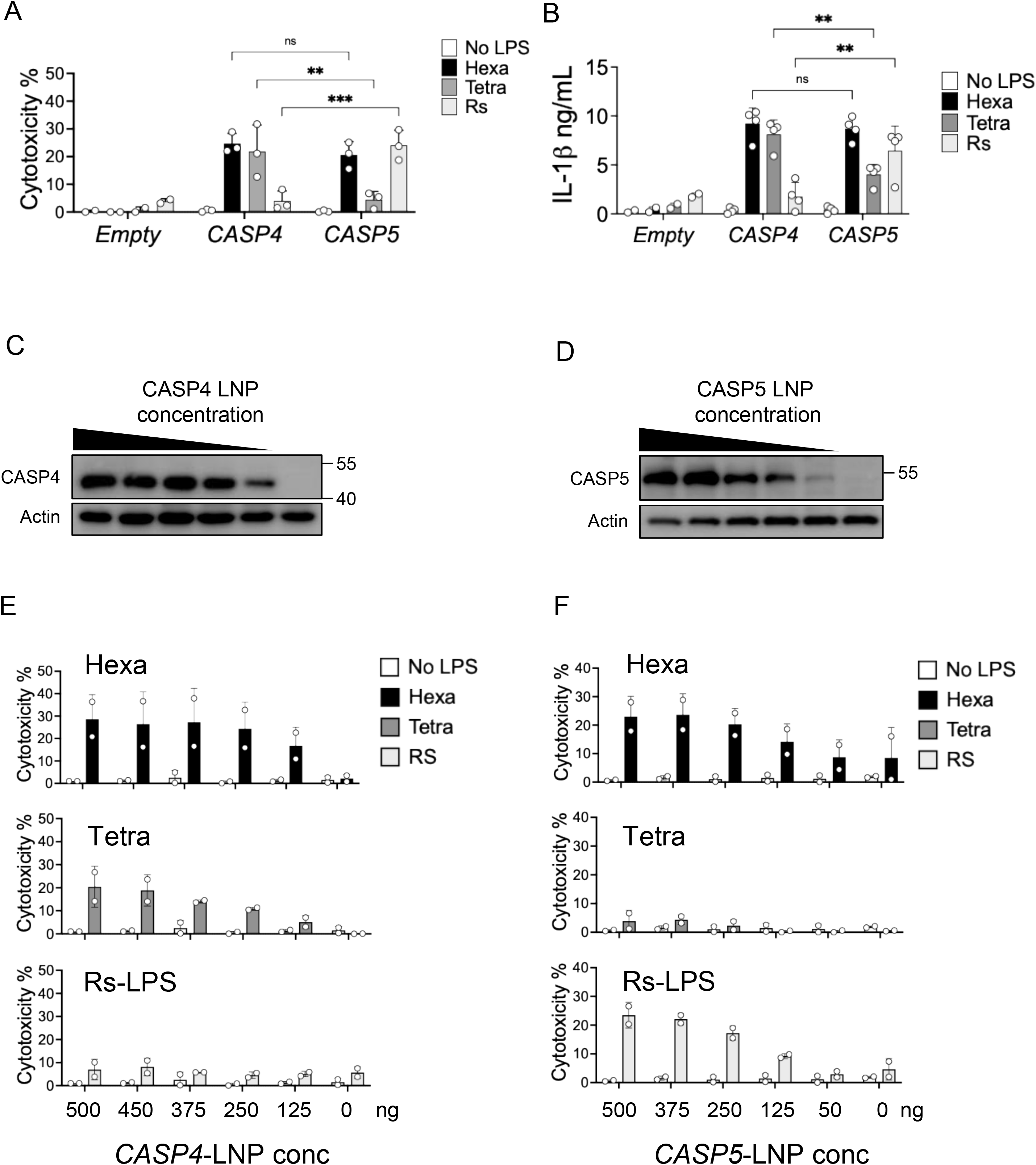
CASP5 and CASP5 exhibit dose-dependent responses to different LPS structures in human macrophages. *CASP4*^-/-^*CASP5*^-/-^ THP-1 macrophages were reconstituted with empty or 0.5µg/E6 cells of LNPs encoding for *CASP4*, or *CASP5* for 48hrs, then primed for 24 hours with PAM3CSK4 (400ng/mL), followed by transfection with indicated LPS structures (2µg/mL): hexa-acylated (Hexa) and tetra-acylated (Tetra) from *Y. pestis*, or Rs-LPS (Rs). Cells were assessed 16 hours later for cytotoxicity by LDH release (A) or IL-1*_β_* release by HEKBlue assay (B). Results show pooled data from at least three independent experiments, with each dot representing the average of three replicates from each experiment (A-B). (C-D) *CASP4*^-/-^*CASP5*^- /-^THP-1 macrophages were reconstituted with the indicated LNPs across a concentration gradient, with caspase expression levels assessed by western blot analysis. (E-F) Cells as in (C-D) were subsequently primed with PAM3CSK4 (400ng/mL) for 4 hours, followed by intracellular stimulation with the indicated LPS structures (2µg/mL), and assessed for cell cytotoxicity 16 hours later. Results are depicted as pooled data from individual experiments, with each dot representing the average of three replicates from each experiment. n.s=not significant, *p≤0.05, **p≤0.01, ***p≤ 0.001 by two-way ANOVA.

### LPS binding is insufficient for non-canonical inflammasome activation

Despite their differential activation by tetra-acylated LPS, Casp11 and CASP4 have previously been shown to bind tetra-acylated LPS^4^. These prior data suggest that non-canonical inflammasome activation is not solely dictated by LPS binding to the caspase. Thus, we next examined whether Casp11, CASP4, and CASP5 interact with hexa-acylated, tetra-acylated, or Rs-LPS. Limited trypsin digestion in the presence of LPS was recently described as an approach to determine the extent of Casp11 interactions with its ligands^28^. This approach is based on the principle that binding of Casp11 to a given ligand, results in conformational changes that alter the accessibility of trypsin cleavage sites within the N-terminal CARD domain, thereby delaying proteolytic cleavage of Casp11 and resulting in a distinct proteolytic digestion pattern that can be visualized by immunoblot analysis. We thus tested the ability of hexa-, tetra-, and Rs-LPS to alter trypsin-mediated caspase digestion. We incubated cell lysates from HEK293T cells stably expressing Casp11, CASP4, or CASP5 with hexa-acylated, tetra-acylated, or Rs-LPS, followed by brief trypsin treatment. Consistent with previous findings, hexa-acylated LPS delayed cleavage of Casp11, as indicated by the persistence of the higher molecular weight band (**Figure S3A**). In contrast, in samples lacking LPS, Casp11 underwent rapid cleavage, resulting in the appearance of a lower molecular weight band.

CASP4 and CASP5 revealed a similar pattern as Casp11 in response to hexa-acylated LPS, which delayed trypsin cleavage of both caspases (**Figure S3B-C**). These findings indicate that the limited trypsin digestion assay can serve as an indicator of interaction between LPS and CASP4 or CASP5. Consistent with prior observations that Casp11, CASP4, and CASP5 can each bind hexa- and tetra-acylated LPS^4^, we found that incubation of Casp11-, CASP4-, or CASP5-containing lysates with hexa- or tetra-acylated LPS led to increased resistance to trypsin-mediated digestion of Casp11, CASP4 and CASP5 (**Figure S3B-C**). Furthermore, Rs-LPS, which has been previously shown to bind Casp11^4^, similarly delayed trypsin-mediation digestion of Casp11, as well as CASP4 and CASP5. These data suggest that both activating and non-activating LPS structures bind to each of these caspases. Nonetheless, LPS binding alone is insufficient to activate these caspases, raising the question of where, downstream of binding, activating and non-activating LPS structures are differentiated^4,28^.

### Activating LPS structures induce formation of non-canonical inflammasome specks

Since non-canonical inflammasome caspase activity correlated with oligomeric speck formation^8^ (**Figure 1**), we hypothesized that only LPS structures capable of activating a cognate caspase would promote the molecular events required for caspase oligomerization and speck assembly, whereas non-activating LPS structures would be unable to do so. We therefore first tested whether tetra-acylated LPS, which weakly activates Casp11, could induce Casp11 speck formation. *Casp11^-/-^* BMDMs expressing Casp11-mCh were stimulated with either hexa-acylated or tetra-acylated LPS and monitored by microscopy for speck formation. Whereas hexa-acylated LPS robustly induced Casp11-mCh specks, tetra-acylated LPS induced similar levels of specks as control-treated cells, indicating that levels of Casp11 activation correlate with speck formation (**Figure 5A-B**).

**Figure 5.**
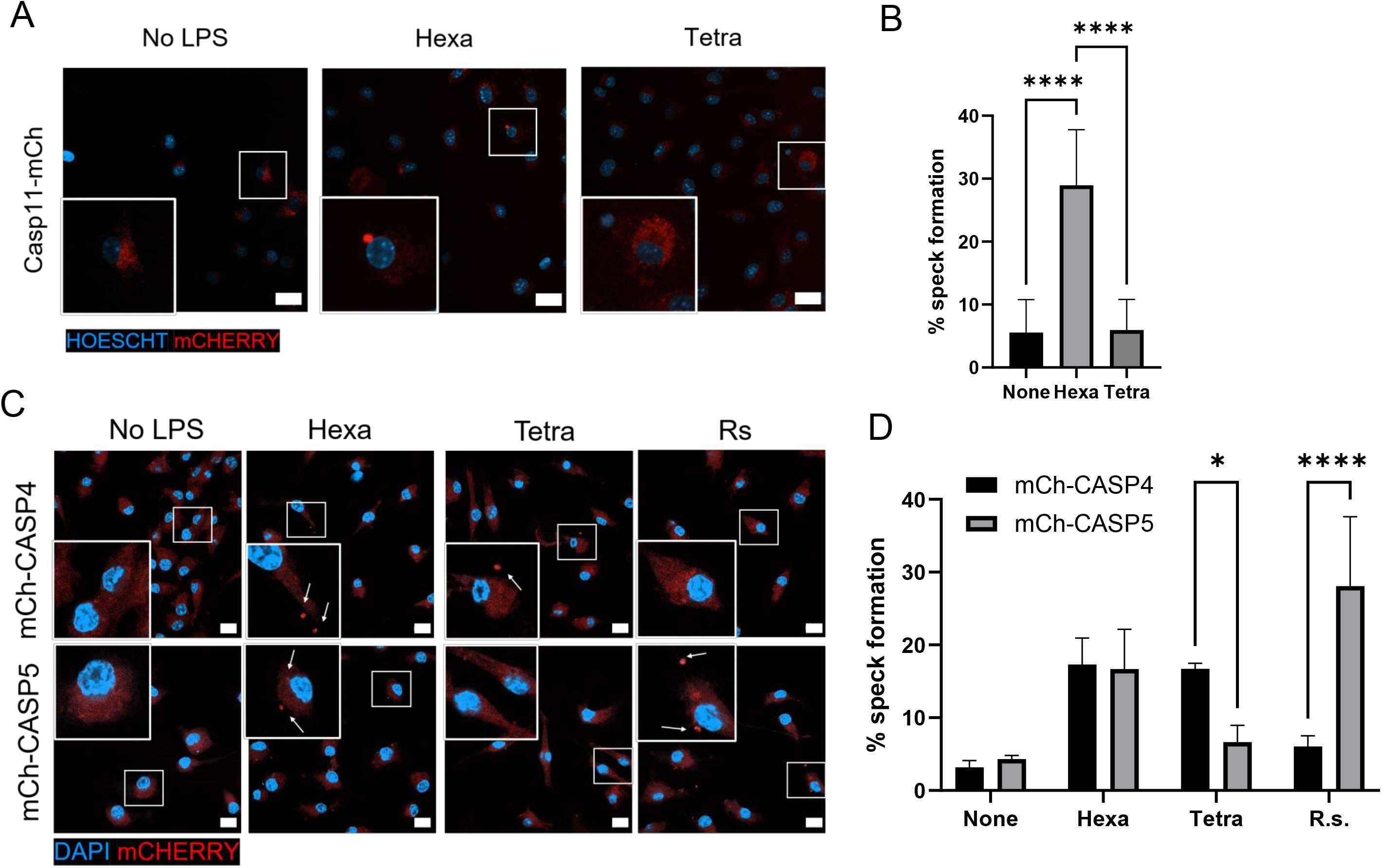
Activating LPS structures induce non-canonical inflammasome speck formation. (A-B) Primary *Casp11^-/-^* BMDMs transduced with lentivirus encoding *Casp11-mCh* were primed with Pam3CSK4 (400ng/mL) for 4 hours, followed by transfection with the indicated LPS structures (2 µg/mL): hexa-acylated (Hexa) or tetra-acylated (Tetra) LPS from *Y. pestis*. 6 hours post-LPS transfection, cells were fixed for confocal microscopy. Nuclei were stained with Hoescht. Images are representative from 3 independent experiments. Scale bar = 20 µm. Speck formation in (B) was quantified as the percentage of Casp11-mCh expressing cells containing at least one speck. Pooled data of 3 biological replicates with 150-200 cells analyzed per condition. ****p<0.0001, One-way ANOVA with Tukey’s multiple-comparison test. (C-D) mCh-CASP4- or mCh-CASP5-expressing iBMDMs were transfected with LPS structural variants (2 µg/mL): hexa-acylated and tetra-acylated LPS from *Y. pestis*, or Rs-LPS. 6 hours post LPS transfection, cells were fixed for confocal microscopy. Nuclei were stained with DAPI. Images are representative from 3 independent experiments. Speck formation in (D) was quantified as the percentage of mCh-CASP4 or mCh-CASP5 expressing cells containing at least one speck. Pooled data of 3 biological replicates with 150-200 cells analyzed per condition. *p<0.05, ****p<0.0001, Two-way ANOVA with Sidak’s multiple-comparison test.

We next assessed speck formation in mCh-CASP4- and mCh-CASP5-expressing iBMDMs in response to activating and non-activating LPS structures. Hexa-acylated LPS induced speck formation in both mCh-CASP4- and mCh-CASP5-expressing cells, consistent with its ability to activate both caspases (**Figure 5C-D**). In contrast, tetra-acylated LPS induced mCh-CASP4 specks but not mCh-CASP5 specks, precisely correlating with its spectrum of activation. Conversely, Rs-LPS, which preferentially activates CASP5, induced robust mCh-CASP5 speck formation, yet minimal mCh-CASP4 specks. Together, these findings demonstrate that LPS-induced speck formation closely correlates with caspase activation. For all three caspases, only activating LPS structures efficiently induce speck formation, whereas non-activating LPS structures did not. Together, these results support a model in which binding is not sole the determinant of activation, and that activating LPS structures specifically mediate caspase oligomerization and SMOC assembly downstream of binding.

### CASP4-mediated responses to tetra-acylated LPS enhance host defense but increase susceptibility to sepsis

Given the differential spectrum of activating and non-activating LPS structures sensed by the murine and human orthologs Casp11 and CASP4, we next considered the possibility that differential sensing of structurally distinct LPS molecules might affect host responses to gram-negative bacterial infections. *Francisella novicida*, a rodent pathogen that naturally expresses tetra-acylated LPS, provided an opportunity to examine the physiologic consequences of differential non-canonical inflammasome activation. In mice, even a small intranasal inoculum of *F. novicida* causes severe pneumonia and bacteremia due to uncontrolled bacterial replication, leading to rapid mortality within one week of infection^29,30^. In contrast, only rarely have human cases of *F. novicida* infection been reported^31^. Intriguingly, THP-1 macrophages lacking CASP4 exhibited higher bacterial burdens than WT THP-1 macrophages following *F. novicida* infection, indicating that CASP4 promotes intracellular bacterial control (**Figure 6A**).

**Figure 6.**
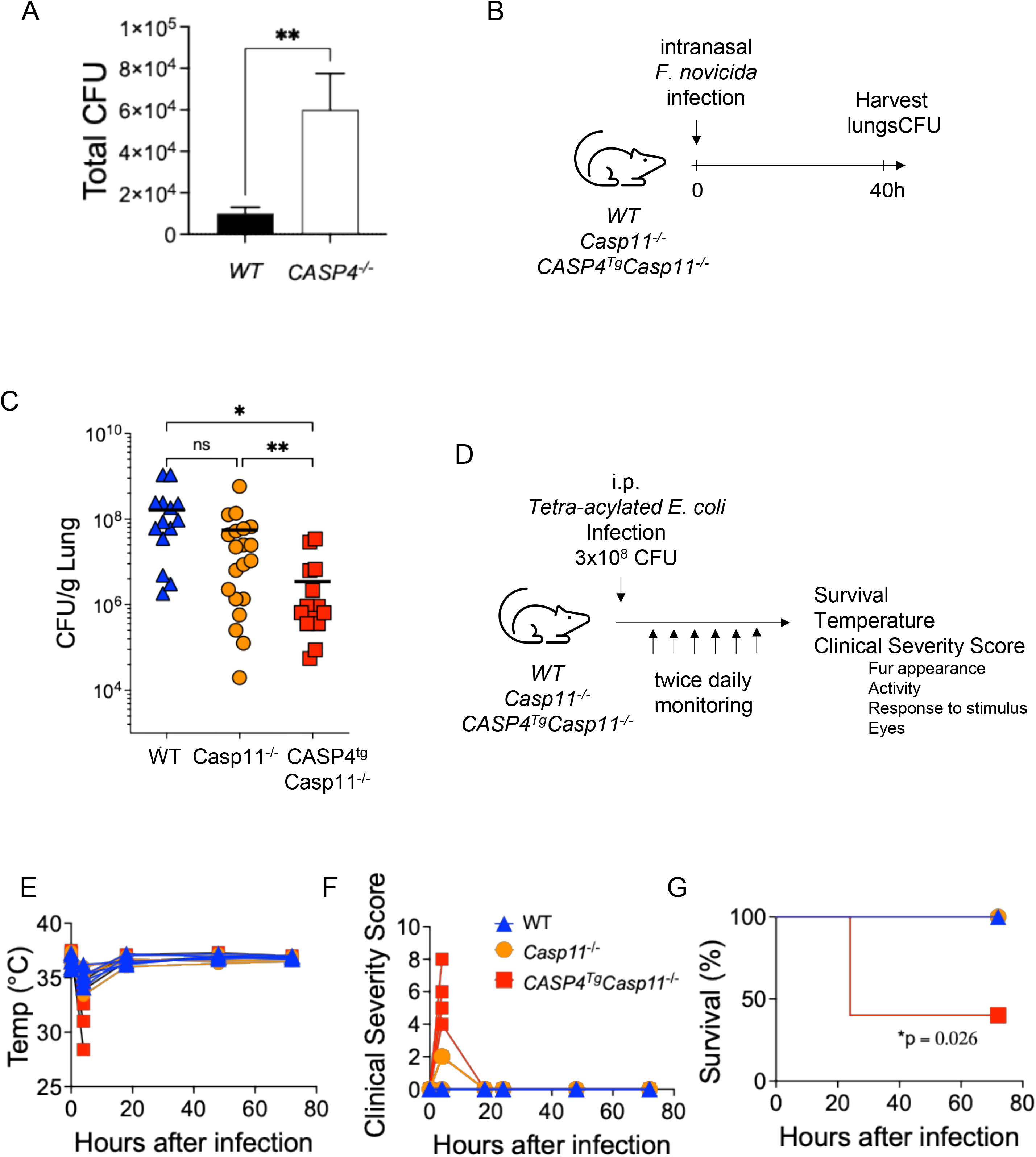
CASP4 responses to tetra-acylated LPS enhance host defense but increase susceptibility to sepsis. (A) PMA-differentiated WT and *CASP4*^-/-^ THP-1 macrophages were infected with *F. novicida* (MOI=20) and assessed for CFUs at 24 hours post-infection. Results are representative of two independent experiments. **p<0.01 by unpaired t-test. (B) Schematic diagram of intranasal *F. novicida* infection. (C) Indicated mice were intranasally infected with 100 colony forming units (CFUs) of *F. novicida.* 40 hours post-infection, lungs were isolated and plated for CFUs. Each point represents an individual mouse. *p<0.05, **p<0.01, ns=not significant by Kruskal-Wallis test. (D) Schematic diagram of tetra-acylated *E. coli* sepsis model. WT, *Casp11^-/-^*, and *Casp4^Tg^Casp11^-/-^*mice were intraperitoneally injected with 3×10^8^ CFU *E. coli* expressing tetra-acylated LPS. Mice were monitored twice daily for changes in body temperature (E), Clinical Severity Score (score of 0-3 in each category with 3 = most severe: fur appearance, response to stimuli, activity, and eyes; max score of 12) (F), and survival (G). Data are representative of four independent experiments with n=5 mice per group. *p=0.026 by Logrank test for survival curve.

We next utilized transgenic mice that express CASP4 but not Casp11 (*CASP4^Tg^Casp11^-/-^* mice) as an *in vivo* model to study how differential sensing of tetra-acylated LPS by CASP4 and Casp11 might impact host defense and systemic inflammation. Consistent with the hypothesis that CASP4-mediated recognition of tetra-acylated LPS enhances antibacterial immunity, *CASP4^Tg^Casp11^-/-^* mice exhibited significantly improved bacterial control following intranasal *F. novicida* infection, resulting in a >2-log reduction in pulmonary bacterial burden compared with WT or *Casp11^-/-^* mice at 40 hours post-infection (hpi) (**Figure 6B-C**). These findings suggest that CASP4-mediated inflammasome responses to tetra-acylated LPS enhance antibacterial host defense, whereas the inability of Casp11 to efficiently respond to tetra-acylated LPS may permit bacterial immune evasion.

Nonetheless, while an inflammatory response to LPS is essential for host defense against invasive bacterial infections, excessive or dysregulated inflammation can drive sepsis and mortality or severe illness. In murine models of gram-negative sepsis, a hyperactivated inflammatory response to LPS is sufficient to cause mortality within 24 hours of high dose LPS exposure, and this response is dependent on Casp11 in mice even when TLR4 is bypassed^1,32,33^. We thus tested whether the increased protection against bacterial infection by might also be linked to increased sensitivity to pathology in response to LPS structures that are not normally sensed by Casp11. To isolate the contribution of LPS structure independent of bacterial virulence factors, we used a genetically engineered nonpathogenic *E. coli* strain expressing tetra-acylated LPS that was specifically designed to induce minimal TLR4 activation and is therefore considered to be endotoxin-free^34,35^. Indeed, despite a high-dose systemic challenge with tetra-acylated *E. coli*, WT and *Casp11^-/-^* mice exhibited little to no clinical response, as determined by low clinical severity scores, minimal temperature changes, and 100% survival, consistent with tetra-acylated LPS being poorly immunostimulatory in mice (**Figure 6D-G**). In contrast, *CASP4^Tg^Casp11^-/-^* mice rapidly developed severe clinical illness, with a substantial decrease in body temperature, high clinical severity scores, and more than 50% mortality within 24 hours. Together, these findings indicate that recognition of under-acylated LPS structures by human CASP4 can enhance antibacterial host defense but also predisposes the host to inflammatory injury and sepsis, revealing a potential evolutionary tradeoff in the expanded range of CASP4-mediated LPS sensing in humans relative to mice.

## DISCUSSION

The cysteine proteases caspase-4 and caspase-5 in humans and caspase-11 in mice are direct sensors of cytosolic LPS and comprise the non-canonical inflammasome. We previously demonstrated that murine Casp11 assembles into supramolecular organizing centers (SMOCs) in response to cytosolic LPS or bacterial infection^8^. Here, we show that human CASP4 and CASP5 also assemble into SMOCs in response to cytosolic LPS or bacterial infection. Like murine Casp11, CASP4 or CASP5 assembly into SMOCs requires their enzymatic activity. These data together suggest that enzymatically coupled SMOC assembly is an evolutionarily conserved feature of non-canonical inflammasome activation.

Prior work from our lab and others showed that the human non-canonical inflammasome respond to a broader repertoire of LPS structures than mice^10,11,15^. In particular, tetra-acylated LPS from *F. novicida* and *Y. pestis* poorly activates the Casp11 inflammasome in mouse macrophages but robustly activates the non-canonical inflammasome in human macrophages^10,11^. However, the individual roles of CASP4 or CASP5 in sensing tetra-acylated LPS or other LPS variants was unclear. Here, we dissect Casp11, CASP4, and CASP5 responses to distinct LPS structures and found that they exhibit differential responses to these structural variants. While hexa-acylated LPS activated all three caspases, tetra-acylated LPS robustly activated CASP4, but not Casp11 or CASP5. To our surprise, penta-acylated Rs-LPS, a well-known antagonist of both TLR4 and Casp11, failed to activate CASP4 but robustly activated CASP5. These data suggest that the two human caspases respond to both overlapping and distinct LPS structures. Future studies will be required to further dissect the structural determinants of LPS molecules that activate CASP4 versus CASP5 and in turn, the unique features of CASP4 and CASP5 that enable them to respond to different LPS structures.

Our data further indicate that LPS binding alone is insufficient for inflammasome activation, as treatment with hexa-, tetra-, or Rs-LPS induced similar levels of resistance to trypsin cleavage in Casp11, CASP4, and CASP5. These data are consistent with prior reports showing that tetra- and Rs-LPS show similar levels of binding to Casp11, despite differential activation^4,28^. We also found that the LPS structures that induced GSDMD processing and cell death also triggered speck formation, while LPS structures that did not cause GSDMD processing or cytotoxicity failed to induce speck formation. These data suggest that speck formation is closely linked with caspase activation by a given LPS variant. While the interaction of these LPS structures with the individual caspases was sufficient to induce conformation changes or impose steric hindrance that limited trypsin digestion, the resulting conformation change is likely insufficient to mediate higher order caspase oligomerization and assembly into active inflammasome complexes. LPS acyl chain number has previously been identified as an important determinant of caspase-4/11 CARD oligomerization^36^. Hexa-acylated LPS promoted the assembly of purified caspase-4/11 CARDs into large octameric complexes, whereas under-acylated LPS structures such as tetra- and Rs-LPS promoted smaller assemblies^36^. In our studies, tetra-acylated and Rs-LPS nevertheless activated CASP4 and CASP5 within macrophages, respectively, suggesting that acyl chain number alone is also insufficient to determine caspase higher order inflammasome assembly. These findings suggest that larger scale structural re-organization, beyond that determined by LPS binding or acyl chain number, is required for functional inflammasome assembly in response to activating LPS structures.

Although murine models have been invaluable tools for dissecting innate immune sensing and non-canonical inflammasome biology, it is likely that fundamental differences in how humans and mice sense bacterial products have hindered our understanding of species-specific host defense or the translation of promising murine sepsis interventions to clinical success. Given that the non-canonical inflammasome response is critical for host defense against bacterial infections, differential sensing of structurally distinct LPS molecules is likely to have direct implications for species-specific host defense and inflammatory responses to LPS *in vivo*. Indeed, we found that mice expressing human CASP4 in lieu of Casp11 better controlled pulmonary infection with tetra-acylated LPS-producing *F. novicida* than WT mice expressing Casp11. However, this enhanced host response to microbial infection was linked to increased susceptibility to sepsis in the setting of non-pathogenic *E. coli* producing tetra-acylated LPS.

Collectively, our data indicate that CASP4 and CASP5 have evolved to detect a broad range of overlapping and distinct LPS structures. Furthermore, like Casp11, both CASP4 and CASP5 assemble into SMOCs that rely on catalytic activity. Critically, LPS-induced oligomerization is selective whereas binding is not. Although activating and non-activating LPS structures show comparable binding to all of the non-canonical inflammasome caspases, only activating structures induce Casp11, CASP4, and CASP5 SMOC assembly. Oligomerization therefore distinguishes activating from non-activating LPS, implicating assembly rather than ligand binding as the step at which the LPS structure is decoded. Consistent with this, the requirement for catalytic activity in oligomerization implies that activating LPS structures induce conformational changes that license oligomerization. How these conformational changes are specified by individual LPS variants, and whether CASP4 and CASP5 utilize distinct mechanisms to assemble into SMOCs remains to be determined. More broadly, our findings define the human noncanonical inflammasome as mechanistically similar to its murine counterpart, but tuned to a broader range of ligands. These findings have implications for our understanding of human-specific responses to gram-negative sepsis and future efforts to target this pathway therapeutically.

## MATERIALS AND METHODS

### Ethics

All animals were handled in compliance with the federal regulations set forth in the Animal Welfare Act (AWA), the recommendations in the Guide for the Care and Use of Laboratory Animals of the National Institutes of Health, and the guidelines of the University of Pennsylvania Institutional Animal Use and Care Committee (IACUC). All protocols used in this study were approved by the IACUC at the University of Pennsylvania (AWA Protocol #804928 and #804523).

### Cell culture

HEK293T cells were purchased from ATCC, grown in complete DMEM supplemented with 10% v/v FBS, 1% L-glutamine, and 1% Penicillin/Streptomycin, and maintained in a 37°C incubator with 5% CO_2_. Murine bone marrow progenitors were harvested from femurs and hip bones of 8– 12-week-old C57BL/6 or Casp11^-/-^ mice and differentiated into BMDMs using 30% L929-conditioned complete RPMI for 7 days as previously described^15^.

*CASP4*^-/-^*CASP5*^-/-^ double-knock out (DKO) THP-1 cells were obtained from Dr. Clare Bryant^22,37^. Cells were maintained in RPMI supplemented with 10% (v/v) FBS, 1% Penicillin/Streptomycin at 37°C incubator with 5% CO_2_. Cells were plated on TC-treated 48-well plates at a concentration of 1 × 10^5^ cells per well for cell cytotoxicity experiments and incubated with 200 µM phorbol 12-myristate 13-acetate (PMA) for 24 hours to allow differentiation into macrophages. Afterwards, PMA-containing media was removed and replaced with media containing the indicated concentrations of LNP for 48 hours. Macrophages were primed with Pam3CSK4 (400 ng/mL; Invivogen) for 4 or 24 hr prior to LPS transfection.

### ER-Hoxb8 conditionally immortalized cell production

Murine myeloid progenitor cells were immortalized according to established protocol^23^. Briefly, bone marrow progenitors were isolated from the femurs and tibias of *Casp11^-/-^*mice. Cells were cultured for three days in RPMI supplemented with 10 ng/mL IL-3 (Peprotech, 213-13), IL-6 (Peprotech, 216-16), and SCF (Peprotech, 250-03) before transduction with MSCVneo-HA-ER-Hoxb8 virus, which was originally created by Dr. David Sykes (Massachusetts General Hospital)^23^. 24 hours after transduction, the cells were replated in R10eg media [RPMI supplemented with 10% v/v FBS, 0.0005 mM beta-estradiol (Sigma Aldrich, E2758), and 10 ng/mL GM-CSF (Peprotech, 315-03)]. 48 hours later, cells were selected for transduction with the addition of 1 mg/mL Geneticin (Thermo Fisher, 10131035). Cells were expanded and passaged in Geneticin-containing R10eg for 2 weeks or until all non-immortalized cells terminally differentiate or die, then maintained in R10eg without Geneticin. To differentiate the myeloid progenitor cells to macrophages, progenitor cells were washed twice with PBS and plated at a density of 1.5 x 10^6^ cells per 10 cm non-TC treated dish in macrophage differentiation media (RPMI supplemented with 20% v/v FBS, 30% L-cell supernatant, 1% penicillin/streptomycin) for 6-7 days.

### Plasmids

pLVV-EF1*_α_*-PGK-Puro lentiviral vector (Genscript) encoding N-terminally-tagged mCherry-CASP4, mCherry-CASP4_C258A, mCherry-CASP5, and mCherry-CASP5_C315A were used for HEK293T transient transfection experiments. pTwist Lenti-SFFV-WPRE lentiviral constructs encoding C-terminally-tagged Casp11-mCherry^8^, CASP4-mCherry, and CASP5-mCherry were utilized for transient transfection experiments. pLEX_307 vectors encoding N-terminally-tagged mCherry-CASP4 and mCherry-CASP5 were used for lentivirus production and transient transfection. Untagged Casp11, CASP4, and CASP5 were encoded on Lenti-SFFV vectors (pTwist) and used for lentiviral production. All constructs were confirmed by sequencing prior to experimentation. Human IL-18 plasmid was purchased from OriGene (catalog# RC202716). Mouse GSDMD plasmid was purchased from Addgene (catalog #80950).

### HEK293T transient transfections

Mammalian expression plasmids were transfected into HEK293T cells using the transfection agent Fugene HD (Promega) at a 3:1 ratio (Fugene:DNA) in Opti-MEM (Gibco), with 125 ng of each DNA plasmid. For cells expressing a single plasmid, 200 ng of the specified plasmid was used. Cells were incubated for 24 hours in a humidified incubator at 37°C and 5% CO_2_ prior to subsequent analysis. For cells incubated with pan-caspase inhibitor zVAD, zVAD (200 μM) was added into appropriate wells at time of transfection and incubated for 24 hours at 37°C and 5% CO_2_.

### Lentiviral production

Lentiviral plasmids encoding caspases (pLEX_307_mCh-CASP4, pLEX_307-mCh-CASP5, pTwist_SFFV_Casp11, pTwist_SFFV_CASP4, and pTwist_SFFV_CASP5) (5 μg each) along with pVSV-G (2.5 μg) and psPAX2 (2.5 μg) packaging plasmids, were transfected into HEK293T cells using Fugene HD (Promega) in Opti-MEM (Gibco) and maintained in a 37°C incubator with 5% CO_2_. After 4 hours, media was changed to complete DMEM (supplemented with 10% v/v FBS, and 1% Penicillin/Streptomycin) and maintained in a 37°C incubator with 5% CO_2_. Viral supernatant were collected at 48 h and 96 h after transfection, combined, and filtered through a 0.45 μm filter, and used for transduction.

### Generating stable mCherry-CASP4- and mCherry-CASP5 -expressing immortalized bone marrow progenitor cells and macrophages

ER-Hoxb8 immortalized *Casp11^-/-^* myeloid progenitor cells were transduced with lentivirus expressing mCh-CASP4 or mCh-CASP5 by incubating viral supernatant with cells in R10eg media. 48 hours later, the media was replaced with R10eg supplemented with 10 µg/mL puromycin (ThermoFisher Scientific cat#A1113803), passaged and expanded for 2 weeks to generate *Casp11^-/-^* progenitor cells stably expressing mCh-CASP4 or mCh-CASP5. Prior to differentiation into macrophages, cells were washed twice with PBS and cultured in RPMI media supplemented with 30% L929 conditioned media, 20% FBS, and 1% penicillin/streptomycin for 6-7 days.

### Generating stable HEK293T cell lines

HEK293T cells were transduced with lentivirus encoding *Casp11*, *CASP4*, or *CASP5* (pTwist-lenti-SFFV vector) and selected with puromycin (1 µg/mL) (ThermoFisher Scientific cat#A1113803), passaged and expanded for 2 weeks with puromycin (1 µg/mL) to generate caspase-stable expressing cells prior to use.

### Trypsin digestion assay

Protocol was adapted from Cao et. al^28^. Cell lysates from HEK293T cells stably expressing Casp11, CASP4, or CASP5 were generated by lysing 5 x 10^6^ cells in 500 µL cell lysis buffer (150 mM NaCl, 25 mM Tris-HCl pH 7.5, 5 mM MgCl_2_, 0.02% digitonin). Equal volumes of cell lysate + LPS (final LPS concentration 1.25 µg/µL) or lysate + buffer were incubated at room temperature for 30mins. 10 µL of each mixture was aliquoted into tubes containing 5 µL lysis buffer. 1 µL trypsin (40 µg/mL) was added, mixed, and incubated for the indicated times. Digestion was stopped by addition of 3.2 uL 6x Laemmli buffer and heating at 95°C for 5 minutes. Samples were then resolved by SDS-PAGE and analyzed by immunoblot (9 μL per well).

### Mice

All animals were housed and bred in specific pathogen-free conditions. WT C57BL/6 mice (Jackson Laboratory), *Casp11^-/-^* mice (Jackson Laboratory)^38^, and *Casp11^-/-^Casp4^Tg^*mice (from Joseph Buxbaum’s laboratory)^39^ were used in this study.

### *In vivo* infections

Intranasal *Francisella novicida* (U112) infection was performed under controlled isoflurane-induced anesthesia. Mice were infected intranasally with 25 μL of a bacterial suspension containing 100 colony forming units (CFU) *Francisella novicida* or PBS vehicle control. At 40 hr post-infection (hpi), mice were euthanized. To measure bacterial CFUs within the lung, lungs were excised and placed into 5 mL sterile water, homogenized using the Miltenyi GentleMACS Dissociator, and cell lysates were then plated onto Tryptic-Soy Agar plates. Bacterial loads were determined by calculating the number of CFU per gram of lung (CFU/g). For tetra-acylated LPS-expressing *E. coli* sepsis model, 3×10^8^ CFU ClearColi^35^ was injected intraperitoneally in a volume of 100 uL per mouse. Animals were monitored twice daily for clinical changes using an adapted Murine Sepsis Severity Score^40^ 0-3 for each category: Appearance (0, Smooth coat; 1, Slightly ruffled fur; 2, Majority of fur on back is ruffled; 3, Piloerection, puffy appearance), Activity (0, Active; 1, Active but avoids standing upright; 2, Active only when provoked; 3, Non-responsive even when provoked 3), Response to stimuli (0, Normal; 1, Slowed response to auditory or touch; 2, No response to auditory, slowed response to touch; 3, No response to touch stimuli), Eyes (0, Open; 1, Not fully open, potential secretions; 2, Half closed, potential secretions; 3, Mostly or completely closed), as well as temperature, and survival.

### *Legionella pneumophila* intracellular infection

*L. pneumophila* serogroup 1 strains were used in all experiments. Lp02-derived (thymidine auxotroph) flagellin-deficient (Δ*flaA*) *L. pneumophila* was cultured on charcoal yeast extract (CYE) agar plates supplemented with thymidine (100 µg/mL) and streptomycin (100 µg/mL) for 48 hr at 37°C prior to infection. The day prior to infection, mCh-CASP4 or mCh-CASP5-expressing iBMDMs were seeded on poly-L-ornithine coated coverglass in 24-well plates (100,000 cells/well) and allowed to adhere overnight in 37°C/5% CO_2_ incubator. On the day of infection, cell media was replaced with OptiMEM without antibiotics and cells were infected with mock or Δ*flaA L. pneumophila* at a multiplicity of infection (MOI) of 10 and incubated for 6 hr in a 37°C/5% CO_2_ incubator. The cells were then fixed with 4% paraformaldehyde for 20 min, washed 3× with PBS, and mounted with DAPI Fluoromount-G Mounting medium (Southern Biotech #0100-20).

### Trichloroacetic acid precipitation and immunoblotting

At the indicated timepoints, whole cell extracts and/or supernatants were harvested and immunoblotted for various proteins. In brief, plates were centrifuged (1200 rpm) to rid supernatants of cell debris. Proteins in supernatants were then precipitated by incubating with 10% 0.61 N trichloroacetic acid (Sigma-Aldrich) for 10 minutes on ice and spun at 13,000 rpm in a 4°C centrifuge. Precipitates were washed with acetone by centrifugation at 13,000 rpm at 4°C and resuspended in 1x protein sample buffer (12.5% 1.5M Tris pH 8.8,10% v/v dithiothreitol, 40mM Tris-HCl pH 6.8, 1.6% w/v SDS, 8% v/v glycerol, 0.08% bromophenol blue). Lysates were harvested in 1x protein sample buffer. Protein samples from lysates and/or supernatants were then prepared for SDS-PAGE by boiling and centrifugation (tabletop max speed; 5 min) before they were run on 12% polyacrylamide gels. Proteins were transferred to polyvinylidene difluoride (Millipore) and immunoblotted with the following primary antibodies: anti-caspase-11 (Novus Biologicals 17D9; 1:1000), anti-GSDMD (Abcam EPR19828; 1:1000), anti-β-actin (Cell Signaling #4967; 1:1000), anti-caspase-4 (Santa Cruz sc-56056; 1:1000), anti-caspase-5 (Cell Signaling D3G4W; 1:1000), anti-IL-18 (MBL PM014; 1:1000). Blots were imaged with the Amersham Imager 680 and chemiluminescence using Pierce RCL Western blotting substrate (Thermo Fisher #32209) or SuperSignal West Femto substrate (Thermo Fisher 34094).

### Confocal microscopy

BMDMs or HEK293T cells were seeded on round, poly-L-ornithine-coated #1.5 circular cover glass (Electron Microscopy Sciences #72230-01) and allowed to adhere overnight. Cells were then transfected with LPS (BMDMs) or indicated DNA constructs (HEK293Ts). At the indicated timepoints, cells were washed with PBS 3× for 5 minutes each, fixed with 4% paraformaldehyde for 10 minutes, and washed with PBS 3× for 5 minutes each. Cells were either stained for 10 min with Hoechst (1 µg/mL), then washed 3× with PBS and mounted with Fluoromount-G Mounting media or mounted with DAPI Fluoromount-G Mounting medium (Southern Biotech #0100-20) and dried overnight in the dark. Slides were imaged on Zeiss LSM980 confocal microscope with Plan-Apochromat 63×/1.4 Oil immersion objective utilizing the 405 nm and 561 nm laser lines for DAPI/Hoescht and mCherry, respectively. Scan mode was frame and bidirectional, and an averaging of 4 was utilized. Program Carl Zeiss ZEN 3.6 was used to process the raw data to construct the images. Processing and filtering settings were kept constant and image intensity was preserved with the raw image scale option in Zen 3.6. Two-dimensional (2D) images were generated. Scale bars were made with the line measurement tool function.

### Image quantification and analysis of caspase speck formation

Each experiment was conducted in three biological replicates. Within each replicate, 150-200 cells per condition were counted per coverslip. A speck was defined as a distinct high-fluorescent perinuclear cluster of mCherry. Speck formation frequency was determined as the percentage of mCherry-(red) cells that contained one or more specks.

### Cytosolic LPS delivery

Following differentiation into mature macrophages, Hoxb8-conditionally immortalized *Casp11^-/-^* BMDMs (iBMDMs) were seeded in TC-treated 24-well (200,000 cells/well) or 48-well plates (100,000 cells/well) in RPMI (supplemented with 10% v/v FBS, 15% L-cell supernatant, 1% penicillin/streptomycin) and left to adhere in a 37°C incubator overnight. On the day of LPS transfection, cells were primed with synthetic bacterial triacylated lipopeptide Pam3CSK4 (400 ng/mL; Invivogen) for 4 hr or 24 hr, or with IFN-γ (100 U/mL) for 24 hr as indicated in the experiment. Prior to LPS transfection, cells were washed once with warmed OptiMEM and resuspended in 300 μL (24 well plate) or 200 uL (48 well plate). LPS from *E. coli (O111:B4)* (Sigma cat #L2630), *Salmonella enterica* serotype Minnesota (Sigma cat # L6261), *R. sphaeroides* (Invivogen cat #prslps), and *Y. pestis*^10^ were packaged into stable complexes using FuGENE HD (0.75μL per sample) (Promega; E2311) and added to cells in Opti-MEM (Gibco). 16 hr following LPS transfection, supernatants were harvested and cells lysed with 1x sample loading buffer (60mM Tris-HCl pH 6.8, 2% SDS, 10% Glycerol, 0.002% bromophenol blue, 5% *_β_*-mercaptoethanol) at a standardized concentration of 6000 cells/μL lysis buffer. To assess for assayed for lactate dehydrogenase (LDH) release as a readout for pyroptosis, cytokine release by ELISA, or underwent TCA precipitation.

### Lactate dehydrogenase (LDH) cytotoxicity assay

At the appropriate timepoints, plates were spun down (1200 rpm) for 3 min to rid supernatants of cell debris. Supernatant (50 μL) was diluted with 50 μL OptiMEM and was then combined with an equivalent volume of LDH reaction buffer/substrate mix (Takara Bio Inc) in a clear-bottom 96-well plate. After 20 min incubation at room temperature (RT), reaction was stopped with 1M HCl and absorbance was read on a spectrophotometer (492 nm) and normalized to mock-transfected cells (FuGENE only; negative control) and cells treated with 1% TritonX-100 (maximal cell lysis; positive control).

### Enzyme-linked immunosorbent assay ELISA

Harvested supernatants were assayed using mouse IL-1β ELISA kit from Biolegend cat #432615 and performed according to manufacturer’s instructions. Absorbance values were read at 450 nm.

### HEK Blue IL-1β reporter assay

To quantify the amount of active IL-1β released, HEK Blue IL-1β reporter cells (Invivogen Cat# hkb-il1bv2) were used according to the manufacturer’s instructions. Briefly, HEK Blue cells were seeded in 96 well plates at 5 x 10^4^ cells/well in DMEM supplemented with 10% FBS and 1% penicillin/streptomycin in a total volume of 180 μL. 20 μL of supernatant from control or treated THP-1 cells were added and the samples incubated overnight at 37°C/5% CO_2_ incubator. 20 μL of the cultured HEK Blue supernatant were mixed with 180 μL Quanti-Blue solution and incubated at room temperature for 1 h before absorbance at 620 nm was read on a BioTek plate reader. Recombinant human IL-1β (Invivogen) was serially diluted and applied to the same plate to generate a standard curve, which was used to calculate the absolute levels of active IL-1β.

### mRNA-LNP design, production, and synthesis

Codon-optimized plasmids encoding fluorescent reporters or hu.CASP4, hu.CASP5, or mu.CASP11 variants were cloned (GenScript). Nucleoside-modified mRNA was produced as previously described using the MEGAscript T7 Transcription Kit (Invitrogen), with complete substitution of uridine with N1-methylpseudouridine (TriLink) and co-transcriptional addition of AG 3’ OMe CleanCap (TriLink)^41^. Following *in vitro* transcription, mRNA was purified with cellulose to remove double-stranded by-products^42^. Sufficient removal of dsRNA was assessed via mRNA transfection of monocyte-derived primary human dendritic cells and subsequent measurement of released IFN-α^43^. Appropriate size and integrity of all mRNAs were evaluated using gel electrophoresis. mRNA was formulated in standard lipid nanoparticle formulations present in the US Food and Drug Administration–approved COVID-19 vaccine using a NanoAssemblr Ignite (Cytiva)^44^. A four-component ethanolic lipid solution (50% SM-102, 10% 1,2-distearoyl-sn-glycero-3-phosphocholine, 38.5% cholesterol, and 1.5% PEG lipid) was microfluidically mixed with mRNA in citrate buffer per manufacturer’s instructions. Buffer was exchanged by dilution with 10% sucrose-PBS (w/v) using 100-kDa centrifugation filtration tubes (Amicon). Appropriate particle size (<100 nm) and polydispersity index (<0.2) were assessed with dynamic light scattering (Malvern).mRNA encapsulation efficiency (>90%) and concentrations were determined using the Quant-iT RiboGreen RNA Assay Kit (Thermo Fisher Scientific) according to the manufacturer’s protocol. mRNA-LNPs were maintained at 4°C for short-term use or stored at an approximate mRNA concentration of 1 mg/mL at −80°C.

### mRNA-LNP mediated delivery

Cells were transfected with LNPs at concentration of 0.5 - 1 μg per 1×10^6^ cells as indicated and incubated for 48 h before LPS transfection.

### Statistical analysis

Data were graphed and analyzed using GraphPad Prism 9. Mean values (± SEM) were compared across triplicate conditions, and p-values were determined using one-sample *t*-test, one-way or two-way ANOVA with Sidak’s multiple comparison as indicated in individual figures.

## Acknowledgments

We thank members of the Shin and Brodsky labs for scientific discussions. We thank Dr. Clare Bryant for providing *CASP4^-/-^CASP5^-/-^* THP-1 cells. We thank the University of Pennsylvania CDB Microscopy Core (RRID SCR_022373) for utilization of the Zeiss LSM 980 confocal microscope. We thank the University Laboratory Animal Resources (ULAR) staff for their assistance with animal husbandry and care during this study. We thank the University of Pennsylvania Summer Undergraduate Internship Program (SUIP) for supporting M.A.E., the University of Pennsylvania Postdoctoral Opportunities in Research and Teaching (PORT) for supporting K.A.W, and the University of Pennsylvania First Exposure to Research in the Biological Sciences (FERBS) for supporting K.I.D.

## Funding

NIH/National Institute of Allergy and Infectious Diseases R01AI194573 (SS, IEB, RKE)

NIH/ National Institute of Allergy and Infectious Diseases R01AI118861 (SS)

NIH/National Heart, Lung, and Blood Institute T32HL007586-40 Training in Pulmonary Immunology (LW)

National Center for Advancing Translational Sciences of the National Institutes of Health K12TR006152 (LW)

National Center for Advancing Translational Sciences of the National Institutes of Health KL2TR001879 (LW)

National Science Foundation Graduate Research Fellowship, DGE-2236662 (SMS)

The Brody Family Medical Trust Fund Fellowship in “Incurable Diseases” of The Philadelphia Foundation (MES)

NIH/ National Institute of Allergy and Infectious Diseases F32AI200195 (MES)

NIH/National Institute of General Medical Sciences Maximizing Investigator’s Research Award (MIRA; Grant No. 1R35GM155239-01) (CYT)

Burroughs Wellcome Fund Investigators in the Pathogenesis of Infectious Diseases Award (Grant No. 1541395) (CYT)

National Institute of General Medical Sciences K12GM081259 (KAW)

## Data and materials availability

All data are available in the main text or the supplementary materials

