## Supplemental Figures for "Human caspase-4 and caspase-5 assemble into non-canonical inflammasomes in response to overlapping and distinct LPS structures"

Supplemental Figure 1.

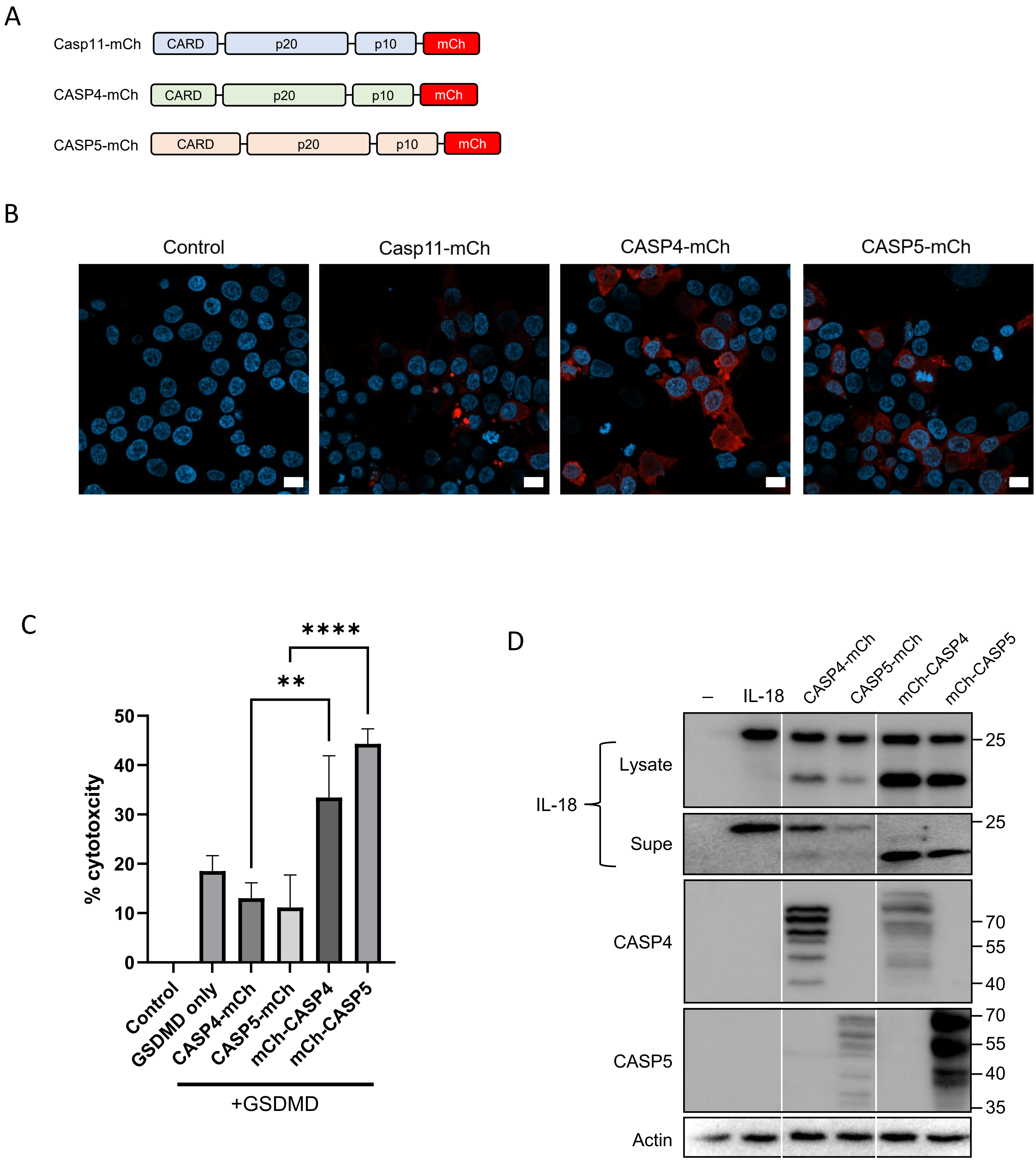

Supplemental Figure 2.

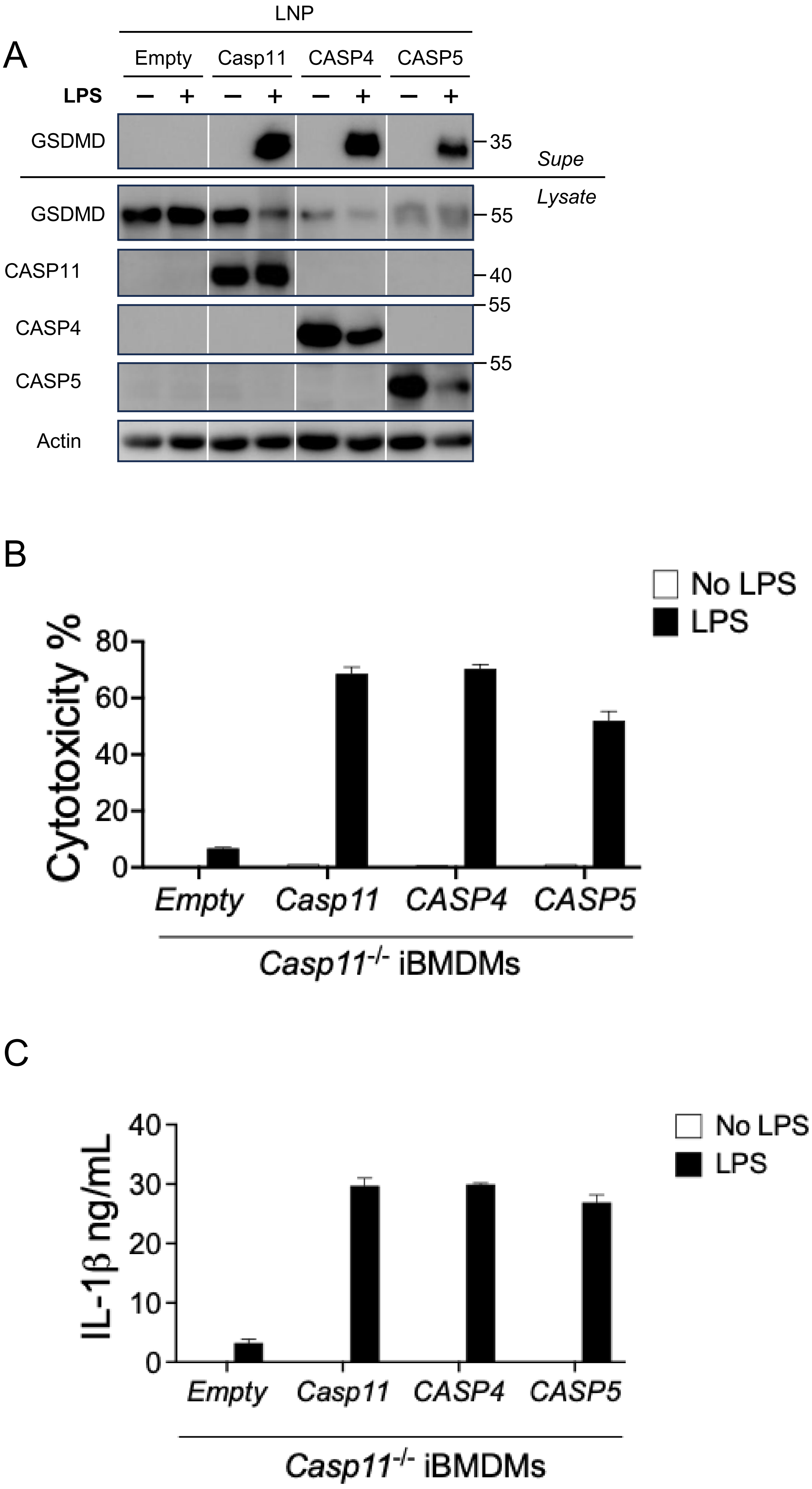

Supplemental Figure 3.

A

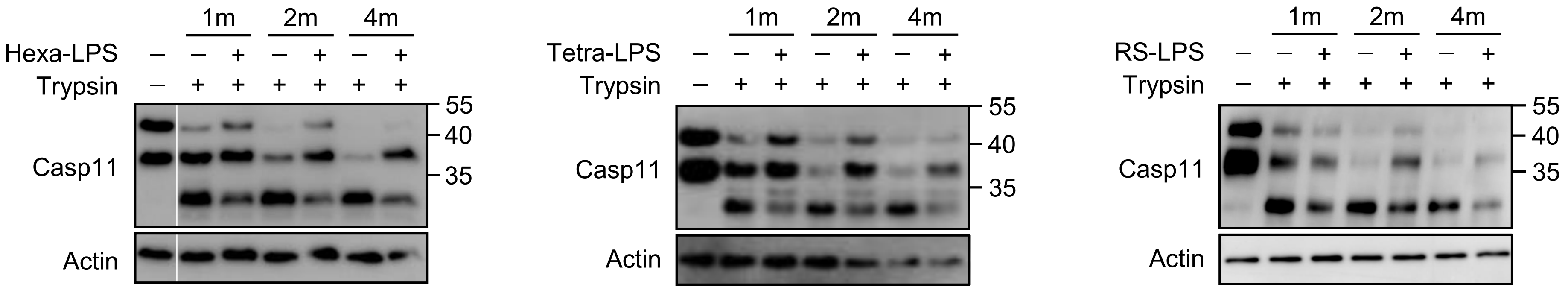


B

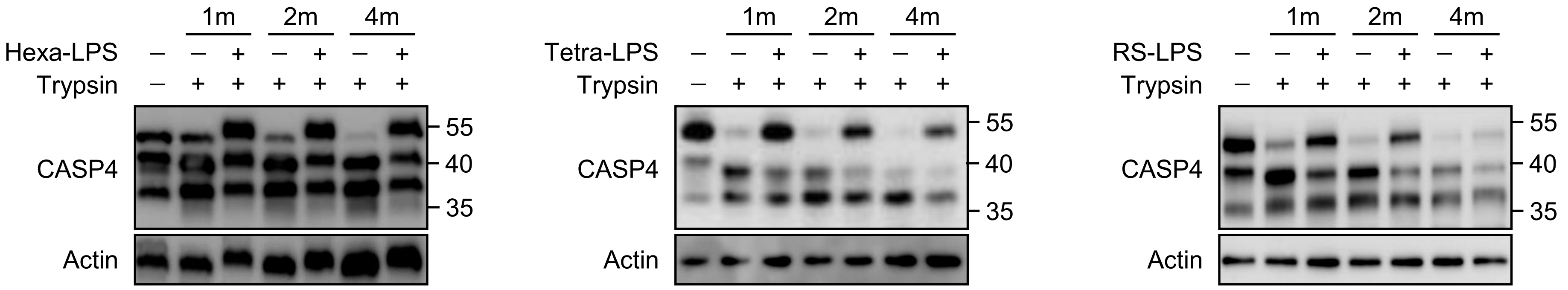


C

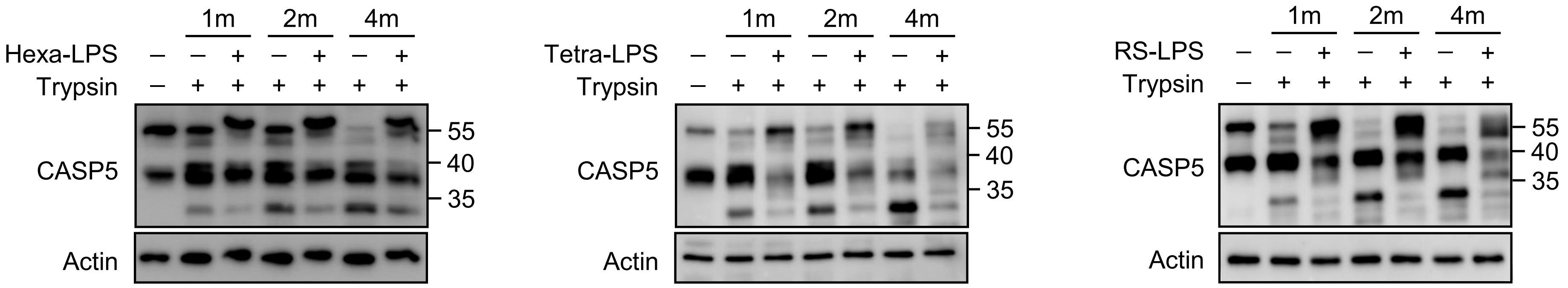

**Supplemental Figure 1. C-terminally tagged CASP4 and CASP5 are poorly functional.** (A) Schematic representation of C-terminal mCherry-tagged Casp11, CASP4, and CASP5. (B) Fluorescence microscopy of HEK293T cells transfected with indicated mCherry- tagged constructs for 24 hours. Nuclei were stained with DAPI. Scale bar = 10 µm. (C) HEK293T cells were transfected with the indicated mCherry-tagged caspase and GSDMD. Cytotoxicity was measured based on lactate dehydrogenase (LDH) release at 24 hr post-transfection. **p<0.01, ****p<0.001 by one-way ANOVA. Shown are pooled data from 3 independent biological replicates. (D) HEK293 T cells co-transfected with plasmids encoding mCherry-tagged CASP4 or CASP5 an IL-18 for 24 hours (125ng each plasmid). Lysates and supernatants were immunoblotted for IL-18, CASP4, CASP5, and β-actin. Representative of at least 3 independent biological replicates.

**Supplemental Figure 2. Human CASP4 and CASP5 are functional in murine macrophages.** *Casp11$^{-/-}$* iBMDMs were reconstituted with 1µg/E6 cells of empty or LNPs encoding for *Casp11*, *CASP4*, or *CASP5* for 48hrs, the primed with PAM3CSK4 (400ng/mL) for 4 hours, followed by intracellular stimulation with hexa-acylated *E. coli* LPS (2µg/mL) for 16hrs and assessed for GSDMD cleavage by immunoblotting for the indicated proteins in the cell lysate or TCA precipitated supernatant (A), cell cytotoxicity by LDH release assay (B), IL-1$\beta$ production by ELISA (C).

**Supplemental Figure 3. Caspase interactions with LPS structures protect from trypsin-mediate digestion**. Lysates from HEK293T cells stably expressing Casp11, CASP4, or CASP5 were incubated with hexa-, tetra-, or Rs-LPS (1.25$\mu$g/uL) for 30mins prior to trypsin (40$\mu$g/mL) exposure for the indicated times prior to immunoblot analysis. Data are representative of at least three independent experiments.
